# Mutations in SARS-CoV-2 Nsp1 are critical determinants of viral pathogenicity in mice

**DOI:** 10.64898/2026.08.14.744970

**Authors:** Nathaniel Jackson, Fuchun Zhou, Anastasija Cupic, Tolga Cagatay, Haiping Hao, Vinay Shivanna, Ruby Escobedo, Kevin Chiem, Chengjin Ye, Lisa Miorin, Beatriz M.A. Fontoura, Adolfo García-Sastre, Alexander Bukreyev, Luis Martinez-Sobrido

## Abstract

Severe Acute Respiratory Coronavirus 2 (SARS-CoV-2) nonstructural protein 1 (Nsp1) dampens the host immune response by shutting off host gene expression, a strategy that has remained evolutionary conserved among a diverse range of coronaviruses (CoVs). Residues important for SARS-CoV-2 Nsp1 mediated host shutoff have been incompletely defined. We have generated and characterized the ability of four Nsp1 mutants to inhibit host gene expression in both plasmid-based overexpression assays and utilized reverse genetic approaches to generate recombinant (r)SARS-CoV-2 expressing each Nsp1 mutant to investigate their impact on viral infection. Infection of K18-hACE2 transgenic mice with the rSARS-CoV-2 Nsp1 mutants resulted in reduced pathogenicity as determined by body weight maintenance and survival, attenuated viral replication in lung and nasal turbinate, distinct immune signatures in lung, and less severe lung pathology in comparison to wild-type (WT) virus-infected mice. Our data suggests amino acid residues in the C-terminal domain, in addition to the linker domain, of SARS-CoV-2 Nsp1 are critical determinants of viral pathogenicity as a result of their role in disrupting host gene expression. The reduced pathogenicity of rSARS-CoV-2 Nsp1 mutants highlights the potential for targeting Nsp1 for rational design of viral inhibitors and development of live-attenuated vaccine strategies as effective prophylactic and therapeutic treatments, respectively, to combat SARS-CoV-2 and possibly other CoV infections.

**IMPORTANCE:** To mitigate the ongoing public health threat posed by Severe Acute Respiratory Coronavirus 2 (SARS-CoV-2) and prepare for future coronavirus (CoV) outbreaks, there is an urgent need for effective prophylactic and therapeutic strategies, including vaccines and antivirals. The nonstructural protein 1 (Nsp1) is a conserved CoV virulence factor that suppresses host gene expression and disrupts immune responses. However, the contribution of specific Nsp1 residues to CoV pathogenesis remains unclear. Here, we identify residues within the C-terminal and linker regions of Nsp1 as critical determinants of SARS-CoV-2 pathogenicity *in vivo*. These findings advance our understanding of CoV host shutoff mechanisms and support Nsp1 as a promising target for the development of live-attenuated vaccines and antiviral therapeutics.

## INTRODUCTION

Coronaviruses (CoVs) had major outbreaks over the past two decades, including epidemics caused by Severe Acute Respiratory Syndrome Coronavirus (SARS-CoV) in 2002-2004 (1, 2); Middle East Respiratory Syndrome Coronavirus (MERS-CoV) in 2012 (3, 4); and the most recent global pandemic caused by Severe Acute Respiratory Syndrome Coronavirus 2 (SARS-CoV-2) which started at the end of 2019 (5, 6). CoVs’ severe disease in humans is in part due to dysregulated immune responses and excessive inflammation (7). This is largely driven by viral antagonism of host innate immune pathways such as retinoic acid–inducible gene I (RIG-I) dependent innate immune responses, with a central role attributed to the nonstructural protein 1 (Nsp1), which is evolutionarily conserved across CoVs (8–10).

SARS-CoV-2 Nsp1 suppresses host gene expression through multiple mechanisms, including binding to the ribosome to block the mRNA entry channel and inhibition of host translation (11, 12) while still allowing transcripts with viral 5’ untranslated regions (UTRs) to be translated (13); interfering with nuclear export of host mRNAs (14, 15); and promoting degradation of host transcripts (10, 16–18). Due to its ability to antagonize host antiviral responses, SARS-CoV-2 and other CoVs Nsp1 represent an attractive target for both antiviral development and rational design of live attenuated vaccines (19, 20).

Structural studies have identified key residues within the C-terminal domain of SARS-CoV-2 Nsp1 that are critical for ribosome interaction and host shutoff activity, and mutation of these residues can restore host gene expression (11, 12). In addition, SARS-CoV-2 Nsp1 contains a linker region that positions the C-terminal domain within the ribosomal cleft to mediate translational inhibition (11). Notably, naturally occurring variants of SARS-CoV-2 have been reported to harbor deletions within this linker region, which may alter C-terminal domain conformation and positioning and impact its interaction with the host ribosome.

In this study, we evaluated four SARS-CoV-2 Nsp1 mutants, including three C-terminal double mutants (K164A/H165A; Y154A/F157A; and R171A/R175A) (11) and one linker region double deletion mutant (ΔK141/ΔS142) derived from a naturally occurring variant (21), to determine their ability to antagonize host gene expression.

We first employed plasmid-based assays to assess the contributions of these mutations in Nsp1 function. However, given the limitations of these systems in recapitulating authentic viral infection, we used reverse genetics approaches to generate recombinant (r)SARS-CoV-2 expressing wild-type (WT) or mutant Nsp1. This strategy enabled us to assess the impact of Nsp1 mutations on viral replication, host shutoff activity, and pathogenicity *in vivo* in the context of viral infection. We found that both the C-terminal and linker region residues are critical determinants of SARS-CoV-2 pathogenicity, with Nsp1 mutants exhibiting attenuation *in vivo*, characterized by reduced viral loads and lung pathology. Furthermore, host responses in Nsp1 mutant-infected animals were distinct from those induced by WT infection, with reduced expression of proinflammatory cytokines and chemokines associated with disease severity, including Monocyte Chemoattractant Protein-1 (MCP-1 or CCL2) (22–24). Collectively, our findings highlight the critical role of Nsp1, and amino acid residues, in SARS-CoV-2 pathogenesis and support the potential of targeting Nsp1 for the development of live-attenuated vaccines and antiviral strategies against SARS-CoV-2.

## RESULTS

### Nsp1 mutants are impaired in their ability to inhibit host gene expression

To evaluate the impact of SARS-CoV-2 Nsp1 mutations on host shutoff activity, three C-terminal double mutants harboring alanine substitutions (K164A/H165A; Y154A/F157A; and R171A/R175A) and one linker region double deletion mutant (ΔK141/ΔS142) were analyzed (**Fig. 1A**). HEK293T cells were co-transfected with pCAGGS plasmids encoding WT or Nsp1 mutants, or empty pCAGGS vector control, together with pCAGGS plasmids expressing GFP and Gaussia luciferase (Gluc) as surrogates for host gene expression. WT Nsp1 served as a positive control for host shutoff, while empty vector represented baseline host gene expression. At 48 h post-transfection, empty vector–transfected cells showed robust GFP expression, whereas WT Nsp1 resulted in a dose-dependent inhibition of GFP expression, with maximal suppression observed with 1000 ng of transfected plasmid (**Fig. 1B**). In contrast, C-terminal mutants (K164A/K165A, Y154A/F157A, and R171A/R175A) exhibited strong GFP expression across all doses, indicating loss of Nsp1-mediated host shutoff activity (**Fig. 1B**). The linker deletion mutant (ΔK141/ΔS142) displayed GFP expression comparable to WT Nsp1, suggesting that this mutation does not impair Nsp1 inhibition of host gene expression (**Fig. 1B**). Quantification of Gluc activity, normalized to empty vector control (set to 100%), yielded results consistent with GFP observations (**Fig. 1C**). All C-terminal mutants (K164A/K165A, Y154A/F157A, and R171A/R175A) showed significantly higher Gluc activity compared to WT Nsp1 at all concentrations tested, whereas ΔK141/ΔS142 behaved similarly to WT Nsp1. Western blot analysis (**Fig. 1D**) further confirmed these findings, with GFP levels correlating with fluorescence (**Fig. 1B**) and Gluc (**Fig. 1C**). Dose dependent expression levels of the C-terminal mutants were readily detected by Western blot whereas WT Nsp1 expression was reduced demonstrating self-inhibitory activity, as observed by decreased protein signal (**Fig. 1D**). Despite being undetectable by Western blot (**Fig. 1D**), the ΔK141/ΔS142 NSp1 mutant retained host shutoff activity comparable to WT Nsp1, indicating functional expression of the mutant protein. Collectively, these results demonstrate that mutations within the C-terminal region of SARS-CoV-2 Nsp1 disrupt its ability to inhibit host gene expression, whereas the deletion ΔK141/ΔS142 within the linker region does not impair this function.

**Figure 1:**
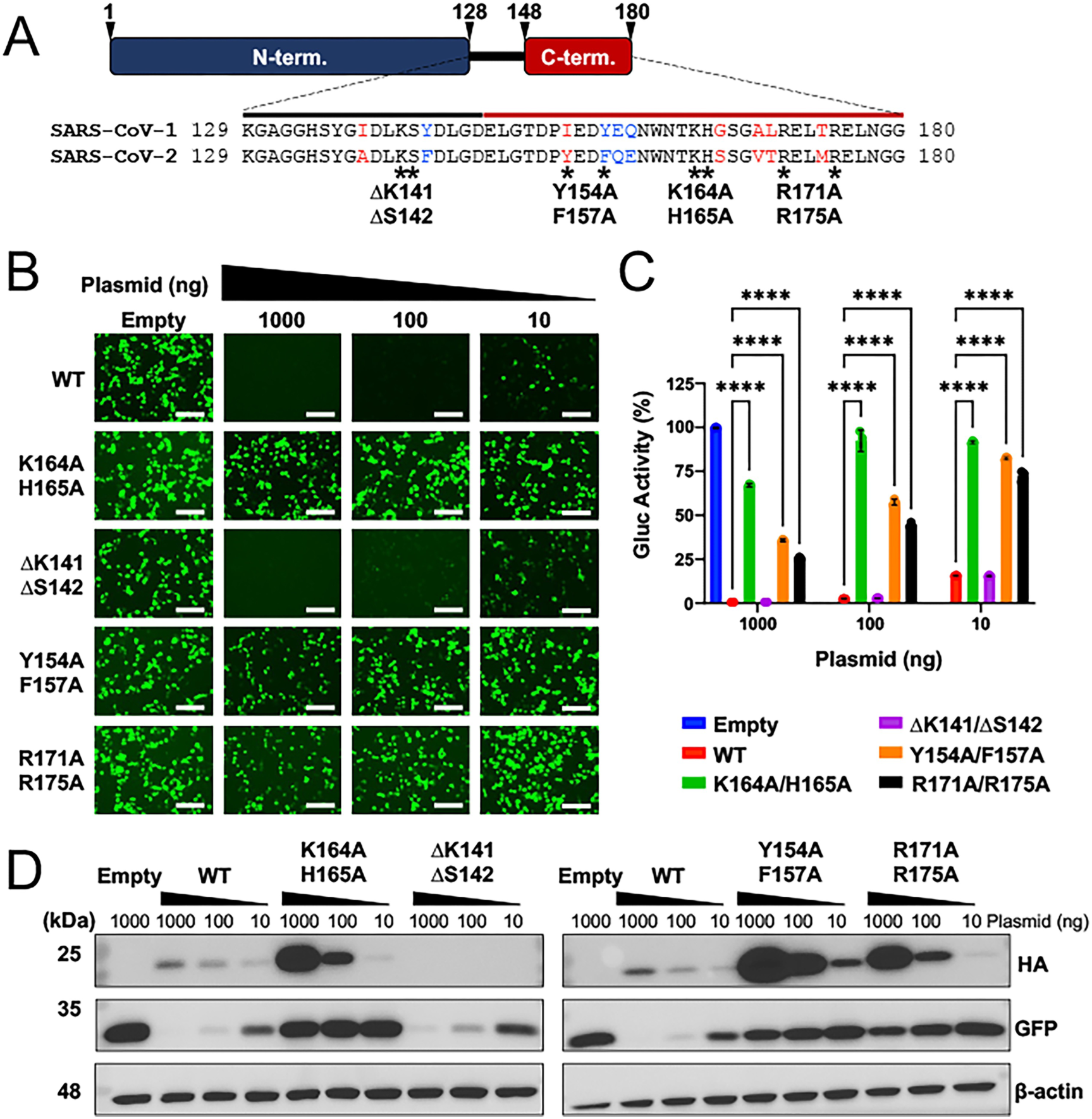
Nsp1 mutants are impaired in their ability to inhibit host gene expression. (**A**) Schematic representation of SARS-CoV-2 Nsp1 mutants. The N-terminal domain is shown in blue, the linker region in black, and the C-terminal domain in red. Sequence alignment of SARS-CoV and SARS-CoV-2 linker and C-terminal domains is shown, with conserved residues in black, similar substitutions highlighted in red, and dissimilar substitutions in blue. Asterisks indicate the positions of introduced mutations. HEK293T cells were co-transfected with (i) 1000, 100, or 10 ng of wild-type (WT) or mutant Nsp1 constructs (all HA-tagged), and (ii) 500 ng each pCAGGS plasmids encoding GFP and Gaussia luciferase (Gluc) reporter plasmids. Empty pCAGGS vector (Empty) was used to normalize total DNA to 2000 ng per transfection and served as a baseline control for host protein expression. GFP expression was visualized at 48 h post-transfection by fluorescence microscopy (**B**) (Scale bar = 100µM), and cell culture supernatants (CCS) were collected to quantify Gluc activity (**C**). Data are presented as the mean of triplicates ± standard deviation (SD). Statistical analysis was performed using two-way ANOVA with Tukey’s multiple-comparison test (****, p < 0.0001). (**D**) Nsp1 protein expression. Cell lysates from cells transfected above were collected and analyzed by Western blot to assess Nsp1 (HA) and GFP expression. β-actin was used as a loading control.

### Nsp1 mutants are impaired in their ability to suppress host protein translation

To specifically assess the ability of SARS-CoV-2 Nsp1 to antagonize host translation, we evaluated the effects of WT and Nsp1 mutants on the translation of an exogenous messenger RNA (mRNA). HEK293T cells were transfected with pCAGGS plasmids encoding WT, mutant Nsp1, or empty pCAGGS vector control for 24 h. Cells were then transfected with *in vitro*–transcribed mRNA encoding firefly luciferase (Fluc). At 10 h post-transfection of Fluc mRNA, cell lysates were collected and Fluc activity was measured as a readout of host translation. Fluc activity was normalized to empty vector control (set to 100%). C-terminal mutants (K164A/K165A, Y154A/F157A, and R171A/R175A) exhibited significantly higher translation levels compared to WT Nsp1, indicating impaired inhibition of host translation (**Fig. 2A**). In contrast, the linker deletion mutant (ΔK141/ΔS142) was not significantly different from WT Nsp1 in inhibiting Fluc expression. Western blot analysis demonstrated robust expression of all the C-terminal mutants relative to WT Nsp1 (**Fig. 2B**). The ΔK141/ΔS142 linker mutant was not detectable by Western blot similar to the results in the plasmid-based host shutoff assay (**Fig. 1D**). These results indicate that mutations within the C-terminal region, and not the double deletion (ΔK141/ΔS142) within the linker region, of SARS-CoV-2 Nsp1 impair its ability to inhibit host translation despite increased Nsp1 expression.

**Figure 2:**
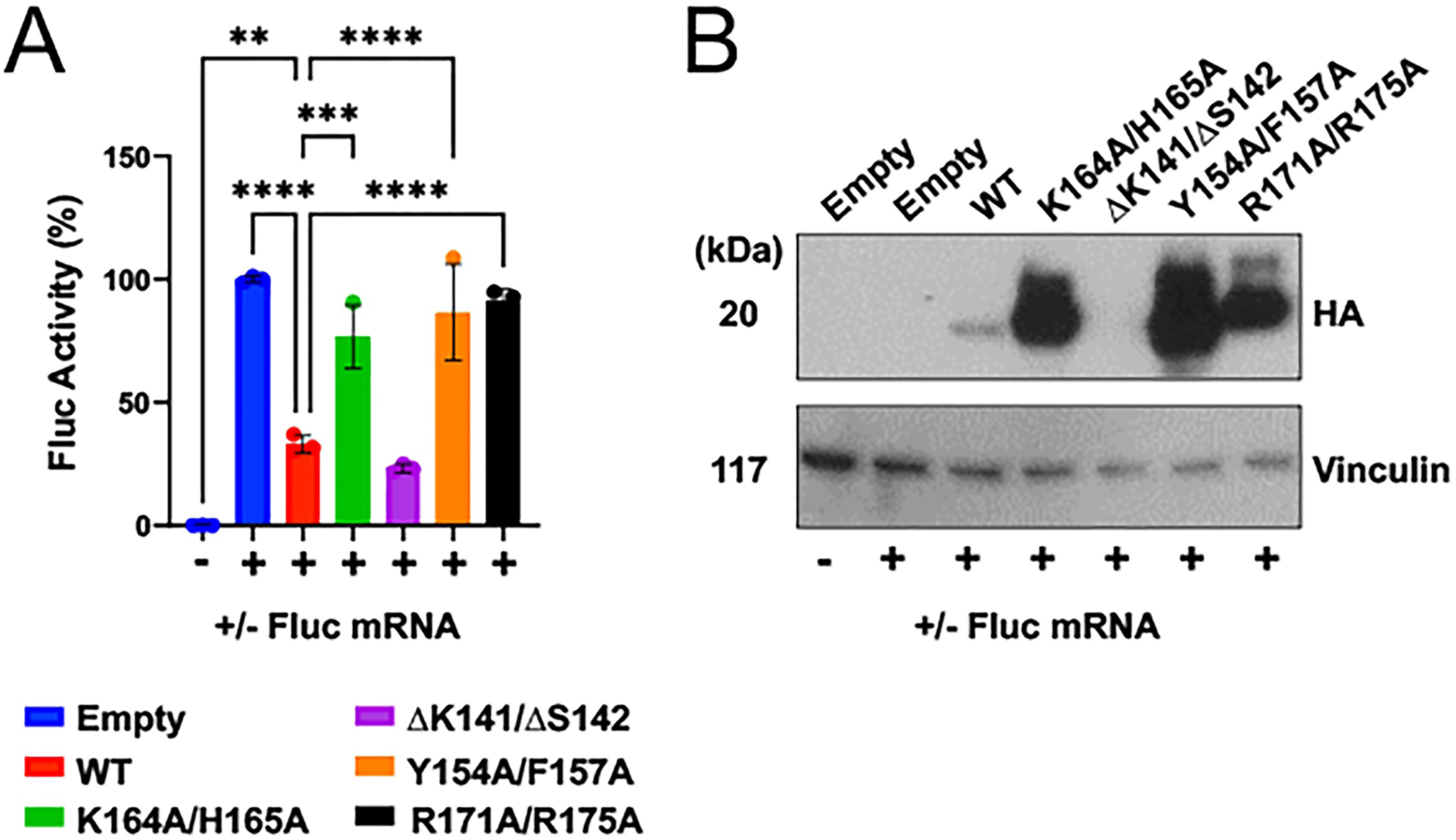
Nsp1 mutants are impaired in their ability to suppress host protein translation. HEK293T cells were transfected with (i) 250 ng of plasmid DNA encoding wild-type (WT), mutant Nsp1 (all HA-tagged), or empty vector control, and 24 h later with (ii) 250 ng of mRNA encoding Firefly luciferase (Fluc). At 10 h post-transfection of Fluc-mRNA, cells were lysed and Fluc activity was measured (**A**). Fluc activity was normalized to the empty vector control and represented as the mean of triplicates ± standard deviation (SD). Statistical analysis was performed using one-way ANOVA with Dunnett’s multiple-comparison test (****, p < 0.0001; ***, p < 0.001; **, p < 0.01). Cell lysates from transfected cells above were analyzed by Western blot to assess Nsp1 expression (HA) (**B**). Vinculin was used as a loading control.

### rSARS-CoV-2 expressing Nsp1 K164A/K165A mutation exhibit reduced viral replication

To evaluate the impact of Nsp1 mutations during authentic viral infection, rSARS-CoV-2 WT and Nsp1 mutant viruses were generated using reverse genetics systems based on a bacterial artificial chromosome (BAC) encoding the full-length SARS-CoV-2 genome (25). The Nsp1 gene was modified to introduce the indicated mutations, and recombinant viruses were rescued by transfection of the WT or mutant BAC plasmids into Vero-E6 cells (19). Cells were subsequently expanded, and successful virus rescue was confirmed by the appearance of cytopathic effect. Rescued viruses were confirmed to harbor the desired Nsp1 mutations by deep sequencing (**Supplemental Fig. 1**). Focus forming assay morphology analysis in Vero-E6 cells (**Fig. 3A, top**) revealed signs of viral attenuation for rSARS-CoV-2 K164A/H165A and Y154A/F157A, as demonstrated by reduced plaque diameters compared to rSARS-CoV-2 WT (**Fig. 3B**). In contrast, plaque analysis in Vero-E6 cells expressing hACE2 and TMPRSS2 (Vero-AT) revealed comparable plaque sizes and morphology across all viruses (**Fig. 3A, bottom**), with the exception of a slight reduction observed for rSARS-CoV-2 R171A/R175A (**Fig. 3C**), demonstrating that plaque size is a cell-dependent effect. Next, we assessed viral replication kinetics in Vero-AT (**Fig. 3D**) and human lung epithelial A549 cells expressing hACE2 (A549-hACE2) (**Fig. 3E**). Most Nsp1 mutants exhibited replication kinetics similar to WT virus in both cell types. However, the C-terminal K164A/K165A mutant virus (green) displayed attenuated replication compared to SARS-CoV-2 WT. While WT and other mutant viruses reached peak titers of approximately 10^8^ TCID₅₀/mL (Vero-AT) and 10^9^ TCID₅₀/mL (A549-hACE2) at 48 h and 72 h post-infection, the K164A/K165A mutant reached ∼10-fold lower viral titers in both cell lines. These results indicate that rSARS-CoV-2 containing the C-terminal K164A/K165A mutation is impaired in viral replication, whereas rSARS-CoV-2 carrying the other C-terminal or the linker Nsp1 mutations are not significantly affected in viral fitness *in vitro*.

**Figure 3:**
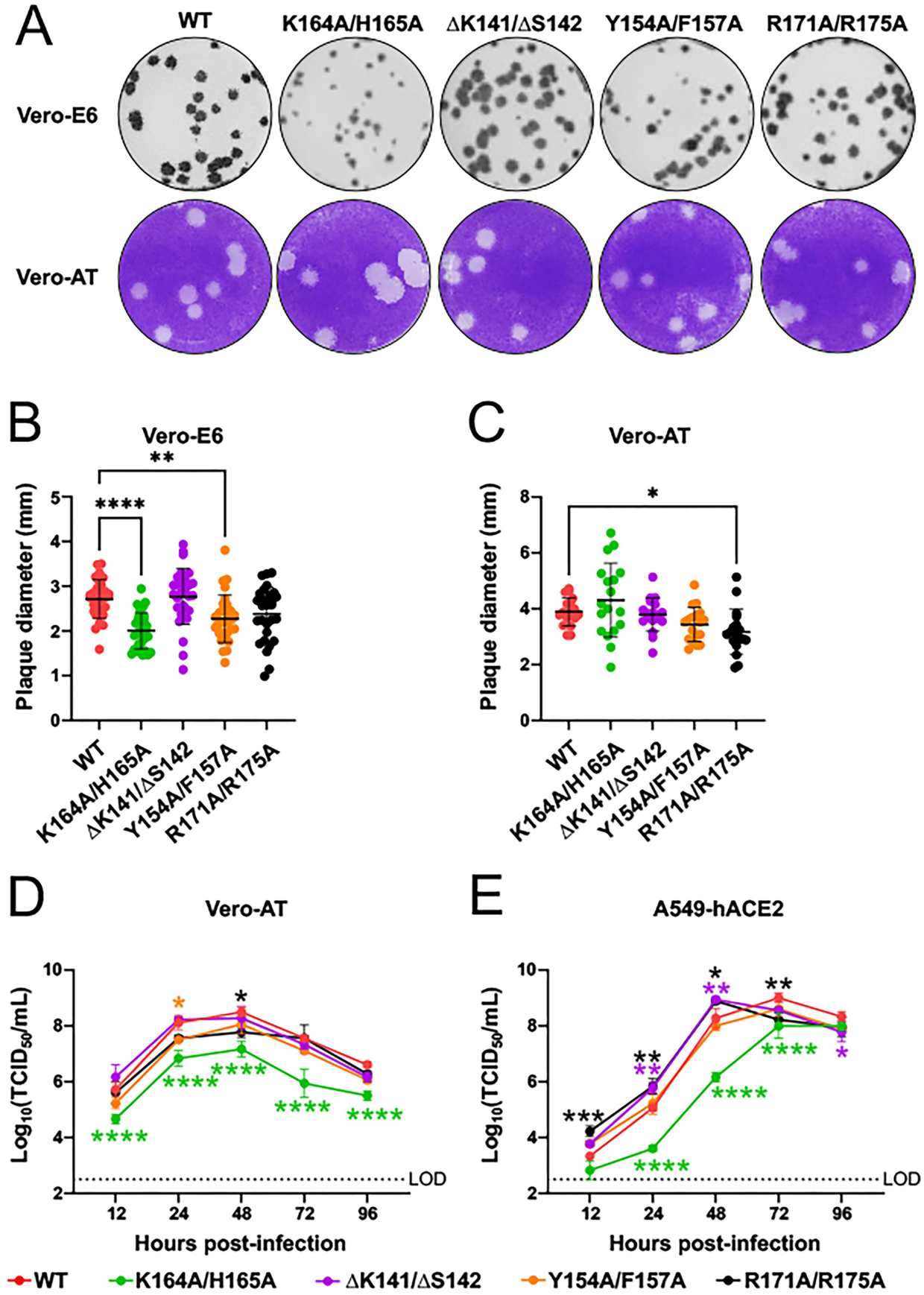
rSARS-CoV-2 expressing Nsp1 mutants exhibit reduced viral replication. Plaque morphology of rSARS-CoV-2 expressing WT or mutant Nsp1 in Vero-E6 (**A, top**) visualized with immunostaining (anti-N) and Vero-AT (**A, bottom**) visualized by crystal violet staining with measured plaque diameter (n = 30 for Vero-E6, n = 18 for Vero-AT) (**B-C**). Data are presented as the mean ± standard deviation (SD) and statistical analysis was performed using one-way ANOVA with Dunnett’s multiple comparisons test (****, p < 0.0001; **, p < 0.01; *, p < 0.05). (**D-E**) Multi-step growth kinetics of rSARS-CoV-2 WT or Nsp1 mutants in Vero-AT (**D**) and A549-hACE2 (**E**) cells infected at MOI of 0.01. Cell culture supernatants were collected at the indicated times post-infection and titrated by TCID₅₀ assay in Vero-AT cells. Data are presented as the mean of triplicates ± standard deviation (SD), and viral titers are expressed as log₁₀ TCID₅₀/mL. LOD, limit of detection. Statistical analysis was performed using two-way ANOVA with Tukey’s multiple-comparison test (****, p < 0.0001; ***, p < 0.001; **, p < 0.01; *, p < 0.05).

### rSARS-CoV-2 expressing Nsp1 mutants exhibit impaired inhibition of host gene expression during infection

To evaluate the inhibition of host gene expression during authentic viral infection, A549-hACE2 cells were mock-infected or infected with rSARS-CoV-2 WT or Nsp1 mutants at a MOI of 0.01. At 48 h post-infection, global host translation was assessed by puromycin incorporation (10). Cells were treated with puromycin for 10 min prior to lysis, and incorporation into nascent polypeptides was analyzed by Western blot (**Fig. 4A**). Mock-infected cells exhibited robust translation, as indicated by a strong puromycin smear, whereas SARS-CoV-2 WT infection resulted in marked inhibition of host translation with minimal puromycin signal **(Fig. 4A**). Among the mutant viruses, the C-terminal mutant K164A/K165A showed the strongest puromycin incorporation, followed by Y154A/F157A, with normalized puromycin-to-β-actin ratios of 16.5 and 3.8, respectively, compared to mock (8.4) and SARS-CoV-2 WT-infected cells (1.5) (**Fig. 4A, bottom**). The linker deletion mutant (ΔK141/ΔS142) exhibited puromycin levels comparable to WT (0.7), while R171A/R175A showed modestly increased translation (2.2) (**Fig. 4A, bottom**). Viral nucleocapsid (N) protein levels were comparable across most infections (**Fig. 4A**), indicating similar levels of viral infection, although K164A/K165A showed reduced N expression, consistent with its attenuated replication (**Figs. 3D-3E**). Robust Nsp1 expression was also detected for all viruses except K164A/K165A, further supporting reduced viral protein accumulation for this mutant due to the lack of efficient viral replication.

**Figure 4:**
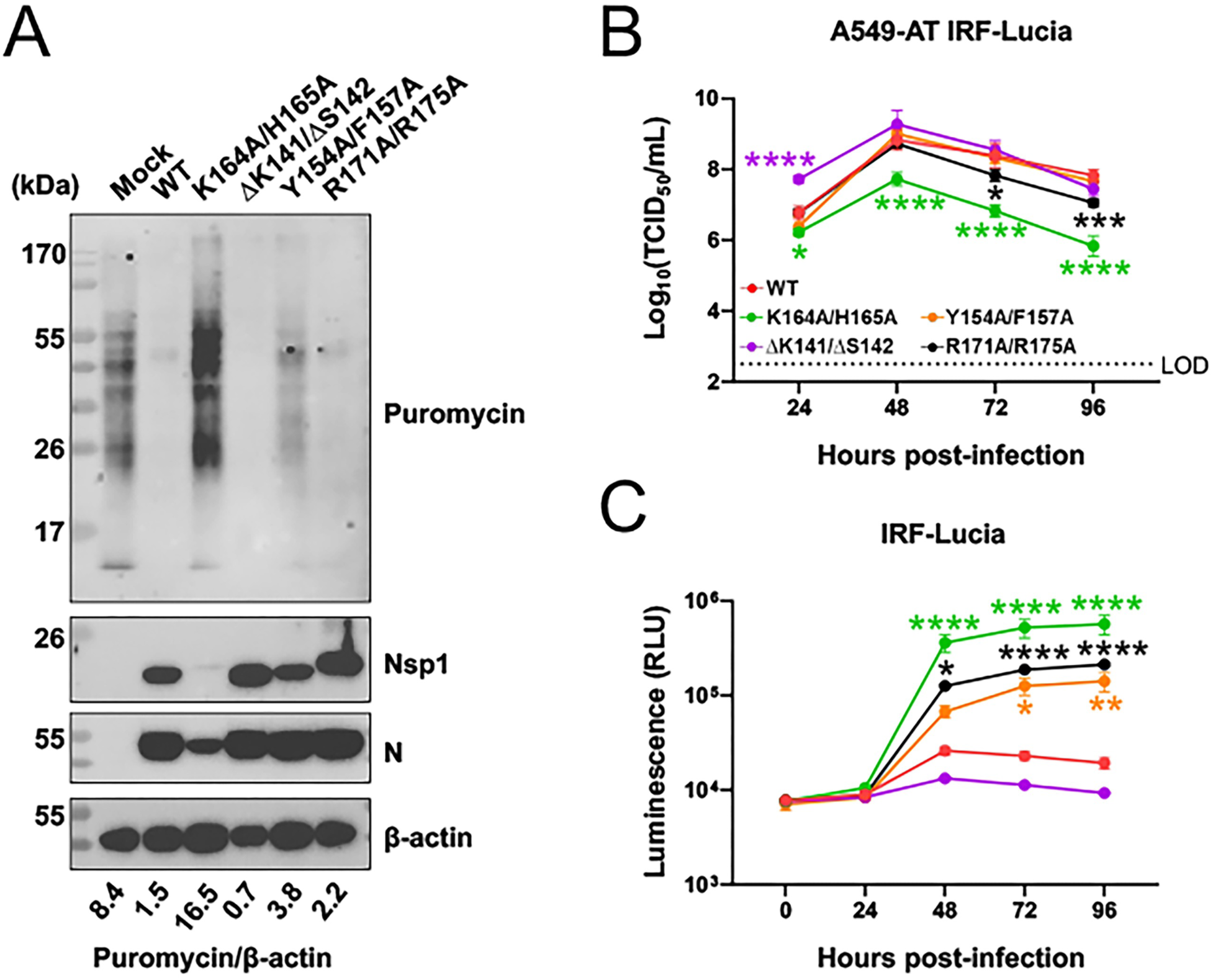
rSARS-CoV-2 expressing Nsp1 mutants exhibit impaired inhibition of host gene expression during viral infection. (**A**) A549-hACE2 cells were infected with rSARS-CoV-2 WT or Nsp1 mutants at MOI of 0.01. At 48 h post-infection, cells were treated with puromycin (10 µg/mL) for 10 min. Cells were lysed and analyzed by Western blot to evaluate host protein synthesis as measured by puromycin incorporation (**A**). Nsp1 and viral N protein expression were also assessed, and β-actin was used as a loading control. Relative translation was quantified as the ratio of puromycin to β-actin based on band intensity (**A, bottom**). (**B**) Multi-step growth kinetics were assessed in A549-AT IRF reporter cells infected with rSARS-CoV-2 WT or Nsp1 mutants. Cell culture supernatants were collected at the indicated times post-infection and titrated by TCID₅₀ assay in Vero-AT cells. Data are presented as the mean of triplicates ± standard deviation (SD), and viral titers are expressed as log₁₀ TCID₅₀/mL. LOD, limit of detection. (**D**) Cell culture supernatants were also assayed for Lucia luciferase activity at the indicated time points. Data are presented as the mean of triplicates ± SD, and raw luminescence values are shown. Statistical analysis was performed using two-way ANOVA with Tukey’s multiple-comparison test (****, p < 0.0001; ***, p < 0.001; **, p < 0.01; *, p < 0.05).

To further assess host responses, A549 reporter cells containing inducible Lucia luciferase reporter under the control of an IRF-responsive promoter, allowing quantification of IRF-dependent transcriptional activity, were modified to express hACE2 and TMPRSS2 to support efficient SARS-CoV-2 infection (A549-AT IRF-Lucia) (**Fig. 4B**). All viruses replicated efficiently in these cells, reaching peak titers of ∼10^8^-10^9^ TCID₅₀/mL at 48 h post-infection, except for K164A/K165A, which exhibited reduced replication (**Fig. 4B**). Luminescence measurements revealed that SARS-CoV-2 WT and the linker mutant (ΔK141/ΔS142) strongly suppressed IRF-dependent transcriptional activity upstream of IFN-induced gene expression (**Fig. 4C**). In contrast, C-terminal mutants induced robust IRF responses, with K164A/K165A, R171A/R175A, and Y154A/F157A showing the highest levels of Lucia luciferase expression (**Fig. 4C**). Altogether, these results demonstrate that C-terminal mutations in Nsp1 impair the ability of SARS-CoV-2 to suppress host responses during viral infection, as evidenced by increased host translation and enhanced IRF-dependent transcriptional activity. Notably, while R171A/R175A showed relatively modest effects on translation inhibition, it induced strong IRF responses, indicating a partial uncoupling of these functions.

### rSARS-CoV-2 expressing Nsp1 mutants are attenuated *in vivo*

To evaluate the impact of Nsp1 mutations on SARS-CoV-2 pathogenesis, K18-hACE2 transgenic mice were intranasally mock-infected or infected with rSARS-CoV-2 WT or Nsp1 mutant viruses using 10⁵ PFU (**Fig. 5A**). Mice were monitored daily for body weight loss (**Fig. 5B**) and survival (**Fig. 5C**). Infection with rSARS-CoV-2 WT was uniformly lethal by day 8 post-infection. Similarly, the rSARS-CoV-2 R171A/R175A mutant closely resembled rSARS-CoV-2 WT infection, with complete mortality by day 8. In contrast, the rSARS-CoV-2 K164A/K165A mutant exhibited partial attenuation, with 25% survival and initial body weight loss (∼10%) followed by recovery to baseline levels by day 8. The rSARS-CoV-2 containing the linker deletion mutation (ΔK141/ΔS142) and the C-terminal mutant Y154A/F157A were the most attenuated, with 75% survival and minimal weight loss, resembling mock-infected mice (**Figs. 5B and 5C**).

**Figure 5:**
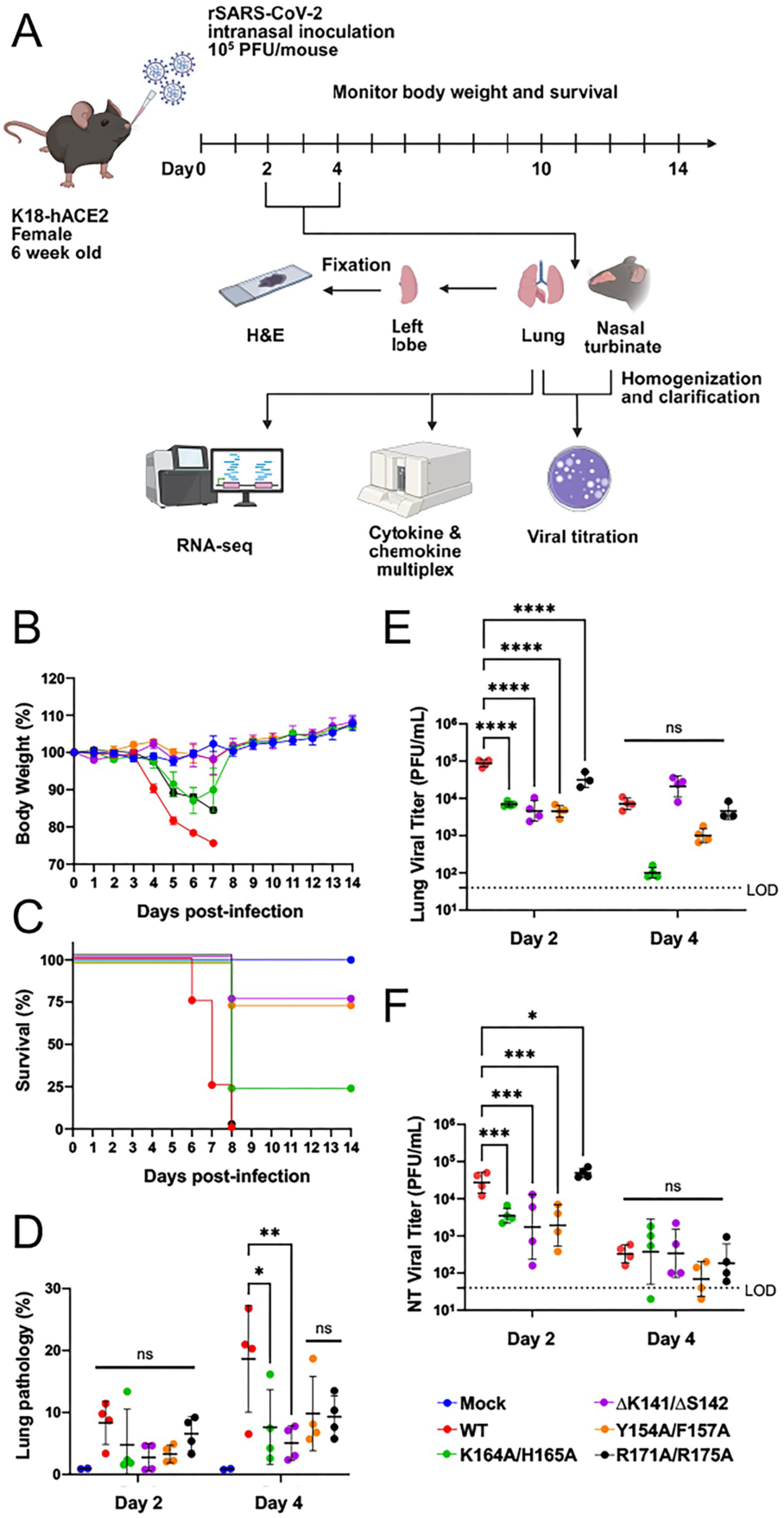
rSARS-CoV-2 expressing Nsp1 mutants are attenuated *in vivo*. **(A)** Schematic representation of mouse experiment and downstream analyses (generated using biorender.com). Six-week-old female K18-hACE2 mice were mock-infected or intranasally infected with rSARS-CoV-2 WT or mutant Nsp1 at a dose of 10⁵ PFU per animal. Animals were monitored for 14 days post-infection for body weight (**B**) and survival (**C**) (n = 4 per group). At days 2 and 4 post-infection, lungs and nasal turbinate (NT) were collected from necropsy groups (n = 4 per group). The left lung lobe was used for histopathological analysis (**D**), and viral titers in the lung (**E**) and nasal turbinate (**F**) were determined by plaque assay in Vero-E6 cells using organ homogenates. Data are presented as the mean ± standard deviation (SD) of animals per group (pathology or viral titers). Statistical analysis was performed using two-way ANOVA with Tukey’s multiple-comparison test (****, p < 0.0001; ***, p < 0.001; **, p < 0.01; *, p < 0.05). LOD, limit of detection; values below the LOD are represented as ½ LOD.

To further assess disease progression, necropsy groups were analyzed at days 2 and 4 post-infection. Lung histopathology at day 4 (**Fig. 5D**) revealed significantly reduced pathology in mice infected with rSARS-CoV-2 K164A/K165A and ΔK141/ΔS142 compared to mice infected with rSARS-CoV-2 WT, consistent with improved survival outcomes (whole-slide lung sections are shown in **Supplemental Fig. 2**). Viral titers in the lungs (**Fig. 5E**) and nasal turbinate (**Fig. 5F**) also correlated closely with survival as lung titers for rSARS-CoV-2 WT and rSARS-CoV-2 R171A/R175A were comparable (∼10⁵ PFU/mL) at day 2 post-infection, whereas the other mutants showed reduced titers (∼10⁴ PFU/mL). By day 4 post-infection, these differences were predominately resolved with the largest reduction seen in rSARS-CoV-2 K164A/K165A (**Fig. 5E**). Nasal turbinate titers followed a similar trend with rSARS-CoV-2 WT and rSARS-CoV-2 R171A/R175A yielding higher viral loads at day 2 post-infection than the other rSARS-CoV-2 Nsp1 mutant viruses. However, differences in viral loads similarly resolved on day 4 post-infection (**Fig. 5F**). Collectively, these results demonstrate that multiple Nsp1 mutations attenuate rSARS-CoV-2 pathogenicity *in vivo*, with the linker deletion mutant (ΔK141/ΔS142) and the C-terminal mutant Y154A/F157A exhibiting the most pronounced attenuation, while other C-terminal mutants, including K164A/K165A, also displaying reduced virulence. In contrast, the C-terminal mutant R171A/R175A retained rSARS-CoV-2 WT-like pathogenicity.

### Lung immune responses are differentially modulated by rSARS-CoV-2 Nsp1 mutants

We next evaluated host immune responses in mice that were mock-infected or infected with rSARS-CoV-2 WT or Nsp1 mutants. Distinct lung cytokine and chemokine expression profiles were observed in mice infected with the rSARS-CoV-2 Nsp1 mutants compared to mice infected with rSARS-CoV-2 WT. At day 2 post-infection (**Figs. 6A-K**), mice infected with the non-attenuated rSARS-CoV-2 R171A/R175A mutant exhibited elevated levels of pro-inflammatory and antiviral cytokines including IL-6, IFN-α, IFN-β, and IFN-γ as well as immune cell-recruiting cytokines and chemokines GM-CSF, CCL2, CCL5, and IP-10 mostly comparable to rSARS-CoV-2 WT. In contrast, expression of these cytokines and chemokines were generally reduced in mice infected with the attenuated rSARS-CoV-2 Nsp1 mutants ΔK141/ΔS142, Y154A/F157A, and K164A/K165A. Most notably, reduced expression levels of IFNs, GM-CSF, CCL2, CCL5, and IP-10 were strongly correlated with survival outcomes. Among these host factors, CCL2 expression was most markedly reduced in mice infected with rSARS-CoV-2 attenuated mutants, strongly correlating with viral attenuation and survival (**Fig. 6I**). By day 4 post-infection, cytokine and chemokine levels had largely converged across groups, with expression patterns approaching rSARS-CoV-2 WT levels, indicating early control of immune responses is critical to prevent severe disease.

**Figure 6:**
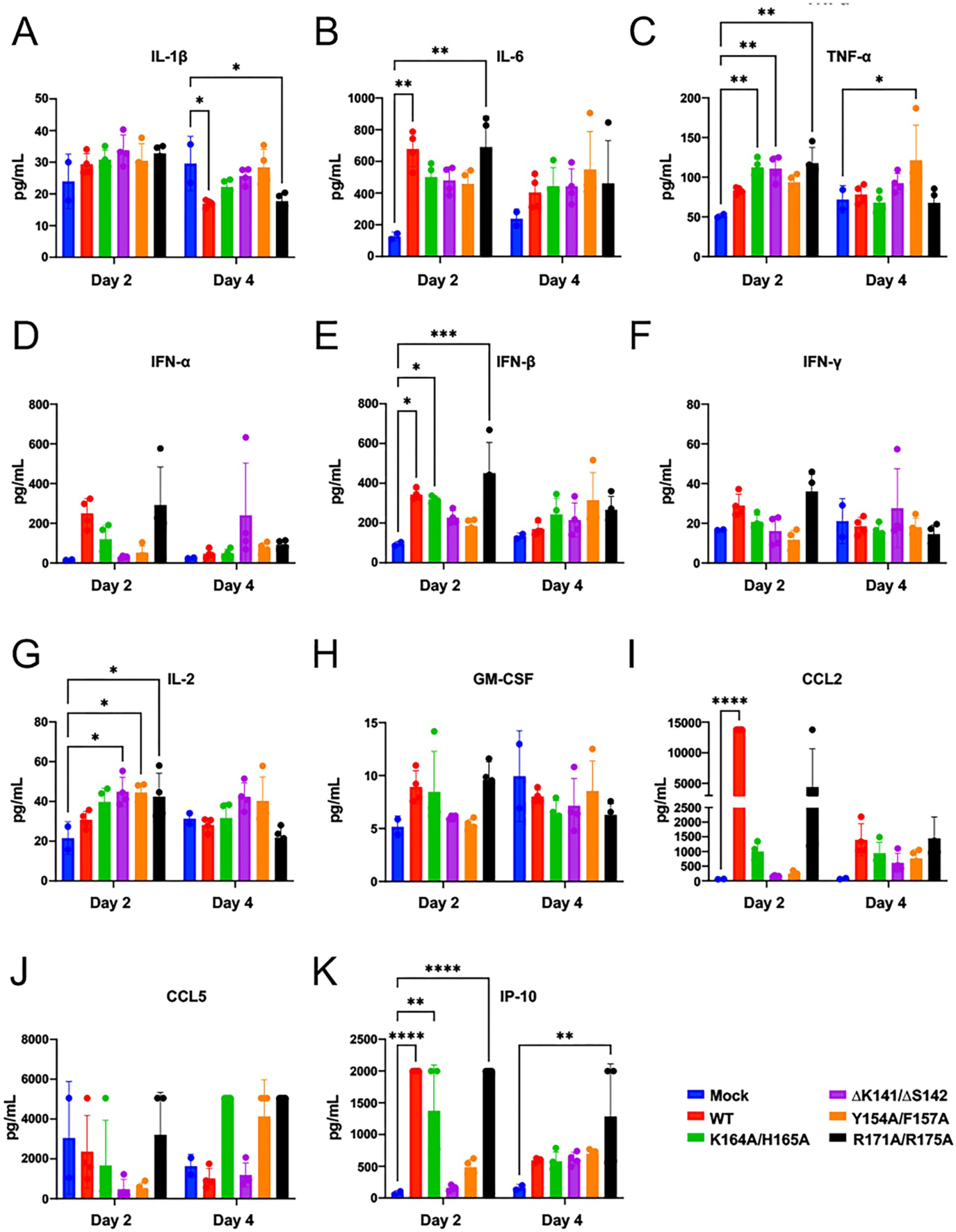
Lung cytokine and chemokine responses are differentially modulated by rSARS-CoV-2 Nsp1 mutants. Lung homogenates for day 2 and day 4 post-infection from mice mock-infected or intranasally infected with rSARS-CoV-2 WT or mutant Nsp1 were analyzed for cytokine and chemokine expression using a Luminex-based multiplex assay. Data are presented as the mean ± standard deviation (SD) of animals per group (n = 4 for virus-infected and n = 2 for mock-infected mice). Statistical analysis was performed using two-way ANOVA with Tukey’s multiple-comparison test (****, p < 0.0001; ***, p < 0.001; **, p < 0.01; *, p < 0.05) comparing to mock-infected animals.

Consistent with proteomic cytokine data, RNA-seq analysis of lung tissues revealed broadly similar global transcriptional profiles, with the greatest differential gene expression relative to mock-infected animals observed in WT-and R171A/R175A-infected mice, whereas attenuated rSARS-CoV-2 mutants exhibited lower overall transcriptional modulation (**Supplemental Fig. 3**). WT-and R171A/R175A-infected mice exhibited elevated transcription of genes associated with inflammation and cytokine signaling, including *Il6*, *Ccl2*, *Ccl7*, and *Cxcl9* (**Fig. 7A**); antigen presentation, including *H2-K1*, *H2-D1*, *B2m*, and *Cd80* (**Fig. 7B**); immune cell markers *Fcgr3* and *Itgam* (**Fig. 7C**); cell death and stress response genes *Bak1*, *Casp8*, *Ripk3*, *Hspa5*, and *Hmox1* (**Fig. 7D**); metabolic response genes *Hk2*, *Eif2s1*, and *Ddit3* (**Fig. 7E**); and tissue remodeling gene *Timp1* (**Fig. 7F**). Collectively, these results demonstrate that Nsp1 mutations alter lung cytokine and chemokine responses *in vivo*, with reduced CCL2 expression emerging as a key correlate of viral attenuation and survival. Furthermore, severe disease was associated with greater transcriptional modulation and increased expression of inflammatory and immune response pathways relative to attenuated rSARS-CoV-2 infection.

**Figure 7:**
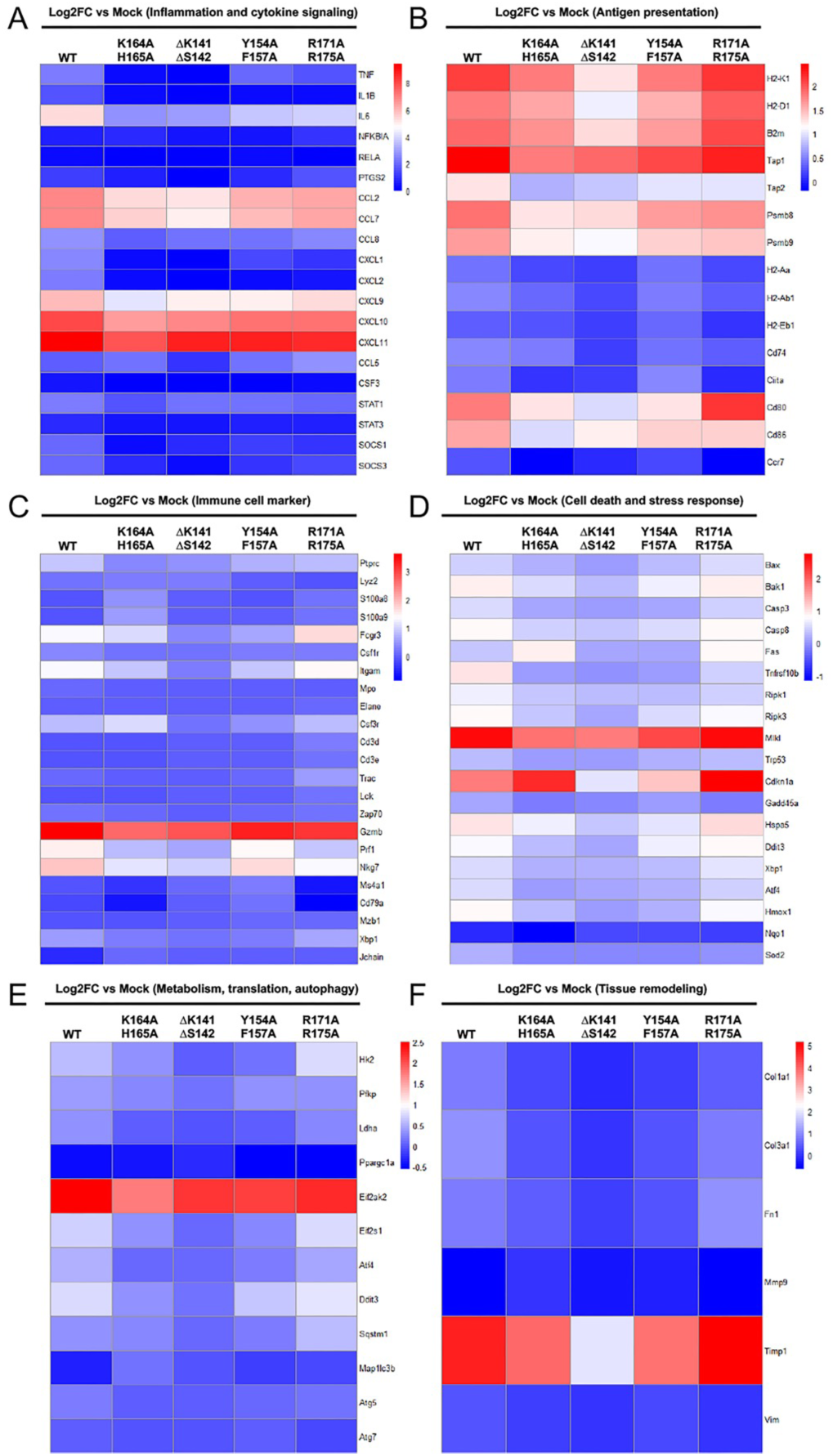
Lung transcriptomic responses are differentially modulated by rSARS-CoV-2 Nsp1 mutants. Lung tissues collected from mice infected with rSARS-CoV-2 WT and Nsp1 mutants at day 4 post-infection were homogenized in TRIzol for RNA isolation and subjected to RNA sequencing (n = 3 per group). Gene expression in WT-and Nsp1 mutant–infected mice were analyzed relative to mock–infected mice and is presented as log₂ fold change (log₂FC). Heatmaps display genes associated with inflammation and cytokine signaling (**A**), antigen presentation (**B**), immune cell markers (**C**), cell death and stress response (**D**), metabolism, translation, and autophagy (**E**), and tissue remodeling (**F**). Downregulation is shown in blue (or expression comparable to mock), intermediate expression in white, and upregulation in red.

## DISCUSSION

We found that the host shutoff activity of SARS-CoV-2 C-terminal Nsp1 mutants was largely consistent with previous overexpression studies (11), with K164A/K165A showing the greatest impairment, followed by Y154A/F157A and R171A/R175A. In contrast, the linker deletion mutant (ΔK141/ΔS142) did not exhibit a discernible defect in host shutoff activity *in vitro* compared to Nsp1 WT. Notably, prior studies evaluating a naturally occurring triple deletion (ΔK141/ΔS142/ΔF143) in Nsp1 reported modest impairment (27, 28), suggesting that shortening of the linker region beyond two-three amino acid residues may be required to more severely disrupt Nsp1 function.

To further dissect the mechanism of SARS-CoV-2 Nsp1-mediated host shutoff, we directly assessed translation using an mRNA-based assay. The resulting phenotypes were consistent with those observed in plasmid DNA-based overexpression host shutoff assays, reinforcing the conclusion that the Nsp1 C-terminal domain predominantly functions at the level of translation via its interaction with the ribosome (11, 12). However, since overexpression systems do not fully recapitulate the context of viral infection we used our previously described BAC-based reverse genetics approaches (25) to generate rSARS-CoV-2 encoding Nsp1 mutants. Using real viral infection, we found that viral replication was largely preserved, with only the C-terminal K164A/K165A mutant exhibiting reduced replication kinetics, consistent with prior reports (10, 19).

Importantly, assessment of host shutoff during authentic infection revealed differences compared to plasmid overexpression systems. While K164A/K165A showed strong impairment of global host translation inhibition and Y154A/F157A displayed a partial defect, R171A/R175A exhibited host shutoff activity comparable to WT, indicating that the effects of Nsp1 mutations are dependent on the context of viral infection. IRF-dependent transcriptional activity was enhanced during viral infection of C-terminal Nsp1 mutants, particularly K164A/K165A and R171A/R175A, suggesting that global translational suppression and selective antagonism of host immune signaling can be uncoupled, as demonstrated by R171A/R175A.

*In vivo* studies revealed a complex relationship between Nsp1 function and pathogenesis. As expected, due to abrogated host shutoff *in vitro*, C-terminal mutants K164A/K165A and Y154A/F157A were attenuated to varying degrees. However, R171A/R175A retained WT-like virulence despite impaired host shutoff activity *in vitro*. Strikingly, the linker deletion mutant ΔK141/ΔS142 exhibited strong attenuation, despite showing WT-like activity in cell-based assays. These findings indicate that Nsp1-mediated virulence is not solely dictated by host shutoff activity measured in standard *in vitro* systems. Analysis of viral titers and lung pathology further supported these observations, with attenuated Nsp1 mutants showing reduced viral burden and decreased lung injury.

Cytokine and chemokine profiling revealed distinct host responses. Infection with rSARS-CoV-2 WT and R171A/R175A were associated with strong induction of pro-inflammatory cytokines and chemokines, whereas attenuated mutants showed reduced expression. Notably, reduced levels of CCL2 expression strongly correlated with survival, consistent with previous studies identifying CCL2 as a key mediator of disease severity (22, 23). Elevated CCL2 has been linked to disease severity in COVD-19 due to excessive recruitment of monocytes and macrophages to the lung (24, 29, 30), contributing to severe acute lung injury, and has therefore been explored as a potential therapeutic target in COVID-19 (31).

Interestingly, despite rSARS-CoV-2 WT retaining robust Nsp1-mediated host shutoff activity *in vitro*, WT-and R171A/R175A-infected mice exhibited broader transcriptional modulation *in vivo* compared to the attenuated Nsp1 mutants and mock-infected mice. Although Nsp1 promotes host mRNA degradation (10, 16–18), RNA-seq analysis at day 4 post-infection likely reflects the cumulative effects of viral pathogenicity, inflammatory amplification, and immune cell infiltration rather than direct host shutoff activity alone. Consistent with this interpretation, WT-and R171A/R175A-infected mice exhibited elevated expression of inflammatory cytokines, antigen presentation genes, immune cell markers, and stress response pathways relative to attenuated Nsp1 mutant viruses.

This might be explained by increased viral replication leading to increased induction of host responses despite a functional Nsp1 in inhibition of host gene expression. These findings suggest that severe disease is associated with greater transcriptional remodeling of the lung environment, potentially driven by enhanced recruitment and activation of inflammatory immune cells (7, 32). In contrast, attenuated SARS-CoV-2 Nsp1 mutants induced reduced inflammatory responses and lower overall transcriptional perturbation, correlating with increased survival.

These results indicate that direct effects of Nsp1-mediated host shutoff may be partially masked at later stages of infection by secondary host immune and pathological responses. Further analysis at earlier time points post-infection will be necessary to better capture the direct transcriptional effects of Nsp1 activity which has been shown to occur early during SARS-CoV-2 infection (i.e. 3 -6 h) (17).

A key unresolved observation is the discordance between *in vitro* and *in vivo* phenotypes, particularly for the linker deletion mutant which demonstrated WT-like activity *in vitro* but attenuated pathogenicity *in vivo* and vice versa for R171A/R175A. Given that Nsp1 exerts its function through interactions with the host proteins such as ribosome (11, 12) and with the mRNA nuclear export factor NXF1 (14, 15), we hypothesize that species-specific differences in host proteins may influence its activity such as differences between human and mice in Nsp1 domains responsible for inhibition of host gene expression.

Overall, our data demonstrate that residues within both the linker and C-terminal regions of Nsp1 are critical determinants of SARS-CoV-2 pathogenicity *in vivo*. We propose that SARS-CoV-2 Nsp1 WT drives dysregulation of the host immune response through its host shutoff activity, whereas Nsp1 mutants deficient in host shutoff permit a more balanced immune response. This is supported by reduced levels of CCL2 in Nsp1 mutant-infected mice, which are associated with more regulated immune cell infiltration and decreased severity of acute lung injury (33). These findings have important implications for the rational design of live attenuated vaccines and are consistent with prior studies demonstrating that Nsp1 mutants (e.g., K164A/K165A) can confer protective immunity in models such as golden Syrian hamsters (20). Furthermore, given that Nsp1 host antagonism is conserved across epidemic CoVs (8–10), our results highlight Nsp1 as a promising target for the development of antiviral strategies and next-generation live attenuated vaccines against present or potential future CoVs outbreaks.

## MATERIALS AND METHODS

### Cell lines

Human embryonic kidney cells (HEK293T; ATCC, CRL-11268), African green monkey kidney epithelial cells (Vero E6; ATCC, CRL-1586), African green monkey kidney epithelial cells expressing hACE2 and TMPRSS2 (Vero-AT; BEI Resources, NR-53821), human lung epithelial cells expressing hACE2 (A549-hACE2; BEI Resources, NR-53821), and human epithelial reporter cells expressing interferon response factors (IRF) with a Lucia luciferase reporter (A549 IRF-Lucia; InvivoGen, a549d-nfis) were used in this study. The A549 IRF-Lucia cells were modified to express hACE2 and TMPRSS2 to generate A549-AT IRF-Lucia cells by lentiviral transduction followed by antibiotic selection to allow SARS-CoV-2 infection. All cells were maintained in Dulbecco’s modified Eagle’s medium (DMEM; Corning) supplemented with 5–10% fetal bovine serum (FBS; VWR) and 100 U/mL penicillin–streptomycin–L-glutamine (Corning) and incubated at 37°C with 5% CO₂.

### Rescue of rSARS-CoV-2 Nsp1 mutants

Recombinant (r)SARS-CoV-2 Nsp1 mutants were rescued from the pBeloBAC11-SARS-CoV-2 infectious clone as previously described (19, 25). Briefly, full-length pBeloBAC11-SARS-CoV-2 plasmids encoding WT or Nsp1 mutants generated by En Passant mutagenesis were transfected into confluent monolayers of Vero-E6 cells using TransIT-LT1 transfection reagent (Mirus Bio) according to the manufacturer’s instructions. At 48 h post-transfection, cells were trypsinized and expanded into larger culture flasks. Viral supernatants were collected upon observation of robust cytopathic effect. To confirm the presence of the desired mutations, viral RNA was extracted from clarified virus stocks using the Direct-zol RNA Microprep kit (Zymo Research) according to the manufacturer’s instructions. Sequencing libraries were prepared for next-generation sequencing and sequenced on an Illumina platform. Reads were aligned to the SARS-CoV-2 reference genome (Wuhan-Hu-1, NC_045512.2), and variant frequencies were determined to assess the presence of mutations in each virus stock.

### Antibodies

#### Host shutoff assay

Primary antibodies used included mouse monoclonal antibodies anti-HA (Cell Signaling Technology, Cat. #2999), anti-GFP (Sigma, Cat. #G1544), and anti-β-actin (Sigma, Cat. #A1978), each at a dilution of 1:1,000.

#### Host translation assay

Primary antibodies used included mouse monoclonal antibodies anti-vinculin (Cell Signaling Technology, Cat. #18799) and anti-HA (Cell Signaling Technology, Cat. #2999), each at a dilution of 1:1,000.

#### Puromycin incorporation assay

Primary antibodies used included mouse monoclonal antibodies anti-puromycin (Sigma, Cat. #MABE343), SARS-CoV cross-reactive anti-nucleocapsid (N) antibody clone 1C7C7 (kindly provided by Dr. Thomas Moran at the Icahn School of Medicine at Mount Sinai), anti-β-actin (Sigma, Cat. #A1978), each at a dilution of 1:1,000, and rabbit polyclonal anti-Nsp1 (kindly provided by Beatriz Fontoura, UT Southwestern).

#### Organ titration immunostaining

SARS-CoV cross-reactive anti-N antibody clone 1C7C7 was used for organ immunostaining at a dilution of 1:1,000.

### Host shutoff assay

#### Transfection, fluorescence microscopy, and luciferase assay

HEK293T cells were seeded at 5 × 10⁵ cells per well in poly-D-lysine–treated (Sigma) 12-well plates in triplicate. Cells were co-transfected with 1000, 100, or 10 ng of plasmid DNA encoding WT or mutant Nsp1 (C-terminal HA tag) in the pCAGGS vector, together with 500 ng of pCAGGS GFP and 500 ng of pCAGGS Gaussia luciferase (Gluc) reporter plasmids using Lipofectamine 3000 (Thermo Fisher Scientific) according to the manufacturer’s instructions. Empty pCAGGS vector was used to normalize total DNA to 2000 ng per transfection and as a control for host gene expression. At 48 h post-transfection, cell monolayers were imaged using an EVOS fluorescence microscope (Thermo Fisher Scientific) to visualize GFP expression. Cell culture supernatants were collected and assayed for Gluc activity using the Pierce Gaussia luciferase assay kit (Thermo Fisher Scientific) according to the manufacturer’s instructions. Luminescence was measured using a Synergy H1 plate reader (Agilent).

#### Western blot analysis

Cells were washed with PBS and lysed using Passive Lysis Buffer (Promega). Triplicate samples were pooled into a single sample. Clarified cell lysates were mixed with SDS-PAGE loading buffer and heated at 90°C for 10 min. Samples were resolved on 12% SDS-PAGE gels and transferred into nitrocellulose membranes (Bio-Rad). Membranes were blocked overnight in 5% nonfat dry milk in PBS (Bio-Rad). Primary antibodies were incubated for 8 h in 5% nonfat dry milk in PBS containing 0.1% Tween-20 (PBST). HRP-conjugated primary antibodies (e.g., anti-HA) were detected directly using chemiluminescent substrate (Thermo Fisher Scientific) and imaged with a ChemiDoc system (Bio-Rad). For non-HRP–conjugated primary antibodies (e.g., GFP and β-actin), HRP-conjugated secondary antibodies (Sigma) were applied for 1 h prior to detection.

### Host translation assay

#### Transfection and luciferase assay

HEK293T cells were seeded in 24-well plates and transfected with 250 ng of pCAGGS plasmids encoding WT or mutant Nsp1 (C-terminal HA tag) using TransIT-LT1. 24 h post-transfection, mRNA encoding firefly luciferase (Fluc) was transfected using Lipofectamine MessengerMAX (Thermo Fisher Scientific), according to the manufacturers’ instructions. At 10 h post-transfection with the mRNA, cells were lysed in Passive Lysis Buffer, and Fluc activity was measured using a luciferase assay system (Promega) according to the manufacturer’s instructions. Fluc activity was normalized to the empty pCAGGS vector control.

#### Western blot analysis

Clarified cell lysates were mixed with Laemmli sample buffer (Bio-Rad), boiled, and resolved on 4–20% gradient SDS-PAGE polyacrylamide gels. Proteins were transferred onto polyvinylidene fluoride (PVDF) membranes (Bio-Rad Laboratories) using the Trans-Blot Turbo Transfer System (Bio-Rad Laboratories). Membranes were blocked in Tris-buffered saline containing 0.1% Tween-20 (TBS-T) supplemented with 5% nonfat dry milk. Primary antibodies were diluted 1:1000 in 3% bovine serum albumin (BSA) in TBS-T.

### Virus titration

#### Standard plaque assay

Vero-E6 or Vero-AT cells were seeded at 5 × 10⁵ cells per well in 6-well plates (1.25 x 10^5^ Vero-E6 per well in 24-well plate for *in vivo* organ titrations) the day prior to infection. Cells were infected with 1 mL (0.25mL for 24-well plate) of 10-fold serial dilutions of virus samples prepared in cell culture medium containing 2% FBS. After 1 h of viral adsorption, the inoculum was removed and replaced with 4 mL (1mL for 24-well plate) of semi-solid overlay consisting of DMEM/F-12 (Gibco) supplemented with 100 U/mL penicillin-streptomycin-L-glutamine (Corning), 0.2% bovine serum albumin (BSA; Sigma), 10 mM HEPES (Gibco), 0.2% NaHCO₃ (Sigma), 0.01% DEAE-dextran (MP Biomedicals), and 1.25% Avicel (IMCD). Cells were incubated for 72 h, fixed with 10% formalin (VWR) overnight, and viral plaques were detected by immunostaining using the SARS-CoV cross-reactive anti-N primary antibody 1C7C7 followed by HRP-based detection with VECTASTAIN reagents (Vector Laboratories) or stained with 0.2% crystal violet (Fisher Scientific) for 5 min prior to plaque visualization. Plaque diameter was measured using ImageJ.

#### TCID_50_ assay

Vero-AT cells were seeded at 2 × 10⁴ cells per well in 96-well plates the day prior to infection. Cells were infected with 100 µL of 10-fold serial dilutions of virus samples prepared in cell culture medium containing 2% FBS. Cells were incubated for 96 h, fixed with 10% formalin overnight and stained with 0.2% crystal violet for 5 min. Titers were calculated using a TCID_50_ calculator excel sheet (Heidelberg University) available at [https://www.klinikum.uni-heidelberg.de/fileadmin/inst_hygiene/molekulare_virologie/Downloads/TCID50_calculator_v2_17-01-20_MB.xlsx].

### Viral growth kinetics

Vero-AT and A549-hACE2 cells were seeded at 5 × 10⁵ cells per well in 6-well plates the day prior to infection. Cells were infected in triplicate with SARS-CoV-2 WT or Nsp1 mutants at a multiplicity of infection (MOI) of 0.01. After 1 h viral adsorption, the inoculum was removed and replaced with culture medium containing 2% FBS. At 12, 24, 48, 72, and 96 h post-infection, cell culture supernatants were collected, and viral titers were determined by TCID₅₀ assay in Vero-AT cells.

### Puromycin incorporation assay

#### Viral infection and puromycin treatment

A549-hACE2 cells were seeded at 5 × 10^5^ cells per well in 6-well plates the day prior to infection. Cells were infected in triplicate with rSARS-CoV-2 expressing WT or Nsp1 mutants at a MOI of 0.01. After 1 h of adsorption, the viral inoculum was removed and replaced with culture medium containing 2% FBS. At 24 h post-infection, cell monolayers were treated with puromycin (10 ug/mL) for 10 min.

#### Western blot analysis

Cells were washed with PBS and lysed using Passive Lysis Buffer. Clarified lysates were mixed with SDS-PAGE loading buffer and heated at 90°C for 10 min. Samples were resolved on 12% SDS-PAGE gels and transferred onto nitrocellulose membranes (Bio-Rad). Membranes were blocked overnight in 5% nonfat dry milk in PBS. Primary antibodies were incubated for 8 h in 5% nonfat dry milk in PBST. HRP-conjugated secondary antibodies (Sigma) were applied for 1 h prior to detection using chemiluminescent substrate and imaged with a ChemiDoc system. Relative translation was determined by quantifying band intensities of the puromycin smear and β-actin using Image Lab software (Bio-Rad). The puromycin signal was normalized to β-actin for each sample.

### IRF luciferase assay

A549-AT IRF-Lucia cells were seeded at 5 × 10⁵ cells per well in 6-well plates the day prior to infection. Cells were infected in triplicate with rSARS-CoV-2 WT or Nsp1 mutants at a MOI of 0.01. After 1 h of adsorption, the viral inoculum was removed and replaced with culture medium containing 2% FBS. At 24, 48, 72, and 96 h post-infection, cell culture supernatants were collected, and viral titers were determined by TCID₅₀ assay in Vero-AT cells. Cell culture supernatants were also used to evaluate Lucia luciferase levels using the Pierce Gaussia luciferase assay kit according to the manufacturer’s instructions. Luminescence was measured using a Synergy H1 plate reader (Agilent).

### Mice experiments

All procedures involving mice were reviewed and approved by the Institutional Animal Care and Use Committee (IACUC) under protocol number 1718MU and conducted in accordance with institutional guidelines for animal care and use. Experiments involving the use of rSARS-CoV-2 WT or Nsp1 mutants were conducted at the ABSL3 facilities at Texas Biomedical Research Institute. Mice were humanely euthanized by Fatal-Plus injection upon reaching humane endpoints, including greater than 25% body weight loss. Survival, weight loss, and viral titers from mice infected with rSARS-CoV-2 WT and K164A/H165A groups have been previously reported (19) and are included here in combination with additional virus-infected groups analyzed from the same study.

#### Viral infection

Six-week-old female K18-hACE2 mice (The Jackson Laboratory) were mock-infected or intranasally infected with rSARS-CoV-2 WT or Nsp1 mutants at a dose of 10⁵ PFU per animal (n = 4 per group). Mice were monitored daily for body weight and survival.

#### Organ collection

At days 2 and 4 post-infection, necropsy groups (n = 4 per group) were humanely euthanized, and lungs and nasal turbinate were collected. The left lung lobe was fixed in 10% formalin and subsequently transferred to 75% ethanol for histopathological processing. The remaining lung tissue and nasal turbinate were homogenized using a tissue homogenizer (Precellys, Bertin Technologies) and clarified for viral load determination and cytokine and chemokine analysis. Additional necropsy groups (n = 3 per group) were euthanized at day 4 post-infection, and whole lungs were excised. Tissues were immersed in RNAlater solution (Thermo Fisher Scientific) and stored overnight at 4°C. Lungs were subsequently washed in PBS and homogenized in 1 mL of TRIzol reagent (Thermo Fisher Scientific). Homogenates were clarified by centrifugation, and total RNA was extracted according to the manufacturer’s instructions (Zymo Research). Purified RNA was used for RNA sequencing.

### Lung histopathology

Left lung lobes were fixed in 10% formalin and subsequently transferred to 75% ethanol. Tissues were paraffin-embedded, sectioned, and stained with hematoxylin and eosin (H&E). Whole-slide images were acquired and analyzed in a blinded manner by a board-certified pathologist. Quantification of percent pathology was performed using HALO image analysis software (Indica Labs).

### Luminex-based multiplex cytokine and chemokine analysis

Cytokine and chemokine levels were quantified using a custom mouse ProcartaPlex multiplex assay (Thermo Fisher Scientific) according to the manufacturer’s instructions. The panel included IL-1β, IL-6, TNF-α, IFN-α, IFN-β, IFN-γ, IL-2, GM-CSF, CCL2, CCL5, and IP-10. Clarified lung homogenates from mock-or virus-infected mice were collected at days 2 and 4 post-infection and used for analysis. Data acquisition and analysis were performed using xPONENT software (Luminex).

### Lung RNA sequencing

RNA sequencing was performed as previously describe (19). Briefly, total RNA was extracted from lung tissues (n = 3 per group) collected at day 4 post-infection using TRIzol reagent (Thermo Fisher Scientific) according to the manufacturer’s instructions. RNA quality was assessed using an Agilent Bioanalyzer. Libraries were prepared using a poly(A) selection-based method and sequenced on an Illumina platform to generate paired-end reads. Sequencing reads were aligned to mm10(GRCm38.p6) and differential gene expression analysis was performed using DESeq2. Differentially expressed genes (DEGs) were identified by comparing WT-and mutant-infected samples to mock virus-infected samples. Log₂ fold change (log₂FC) values relative to mock-infected mice were used for downstream visualization, and heatmaps were generated in R.

### Statistical analysis

All statistical analyses were performed using GraphPad Prism (version 10.6.0). *In vitro* experiments were conducted in biological triplicates and data are presented as mean ± standard deviation (SD). For comparisons involving multiple groups across different Nsp1 concentrations (plasmid-based assays), multiple time points (viral titrations and cytokine/chemokine multiplex), two-way analysis of variance (ANOVA) followed by Tukey’s multiple-comparison test was used. For comparisons involving a single Nsp1 concentration across multiple groups or plaque diameter of rSARS-CoV-2, one-way ANOVA followed by Dunnett’s multiple-comparison test was applied. Statistical significance was defined as ****, P < 0.0001; ***, P < 0.001; **, P < 0.01; *, P < 0.05; and ns, not significant. The limit of detection (LOD) is indicated where applicable, and values below the LOD were assigned a value of one-half the LOD for analysis.

## ACKNOWLEDGMENTS

We thank Dr. Thomas Moran for the kind gift of the SARS-CoV cross-reactive anti-nucleocapsid (N) antibody (clone 1C7C7). We thank Dr. Reagan Meredith for assistance with R scripting for RNA-sequencing data analysis. This work was partly funded by the Center for Antiviral Medicines & Pandemic Preparedness (CAMPP), an NIAID supported AViDD (Antiviral Drug Discovery) Center (U19AI171443, to AG-S, LM-S and BF), and by NIAID grant R01AI184975 (to AG-S and BF).

---

**Supplemental Figure 1:**
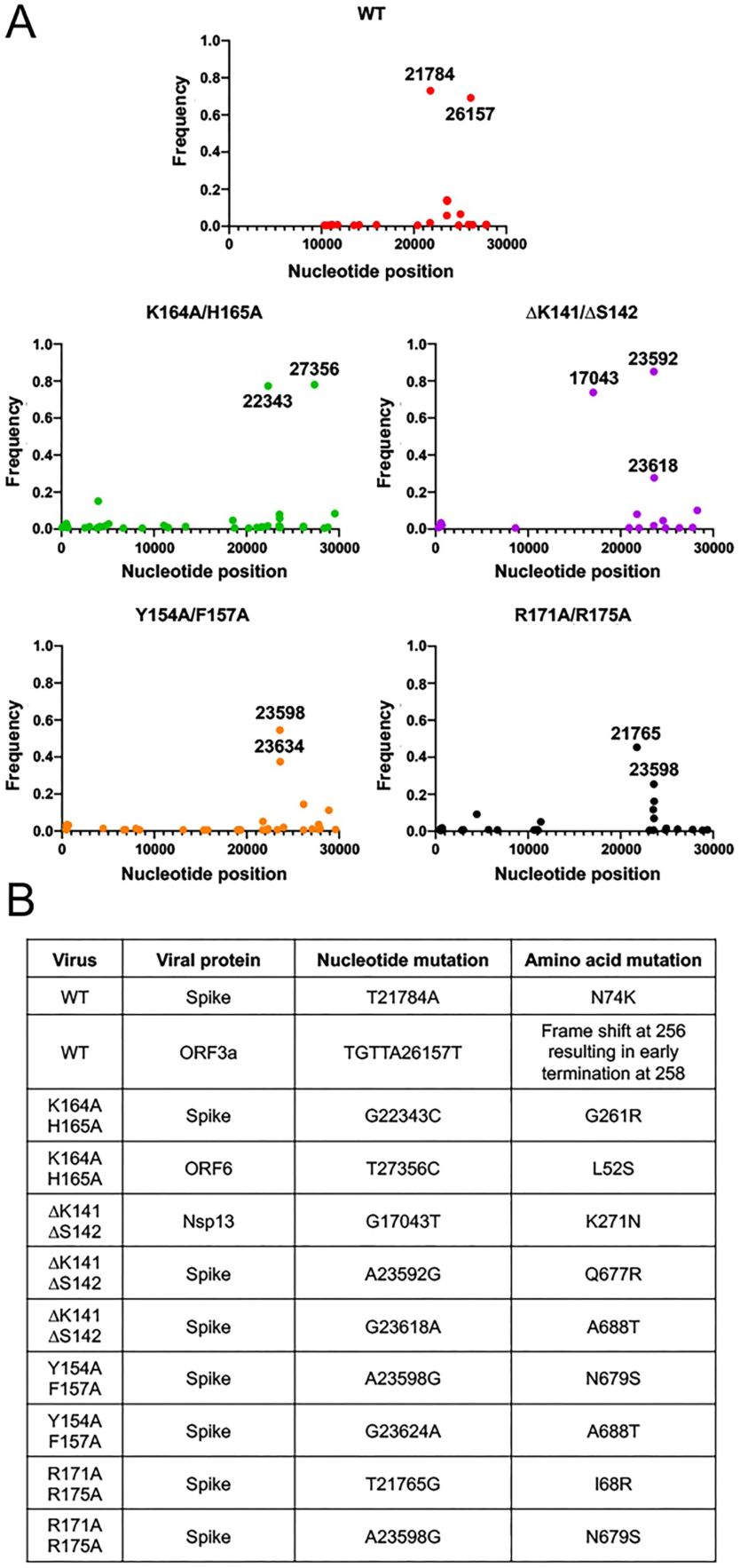
Deep sequencing of rSARS-CoV-2 Nsp1 mutants. Viral RNA was isolated from virus stocks and subjected to deep sequencing to confirm the presence of desired Nsp1 mutations. Reads were aligned to the WT reference sequence or to the sequences containing the respective Nsp1 mutations. The frequency of non-consensus mutations across the viral genome is shown for each virus (**A**). Variants with a frequency greater than 20% are summarized in the table (**B**), along with their corresponding viral proteins.

**Supplemental Figure 2:**
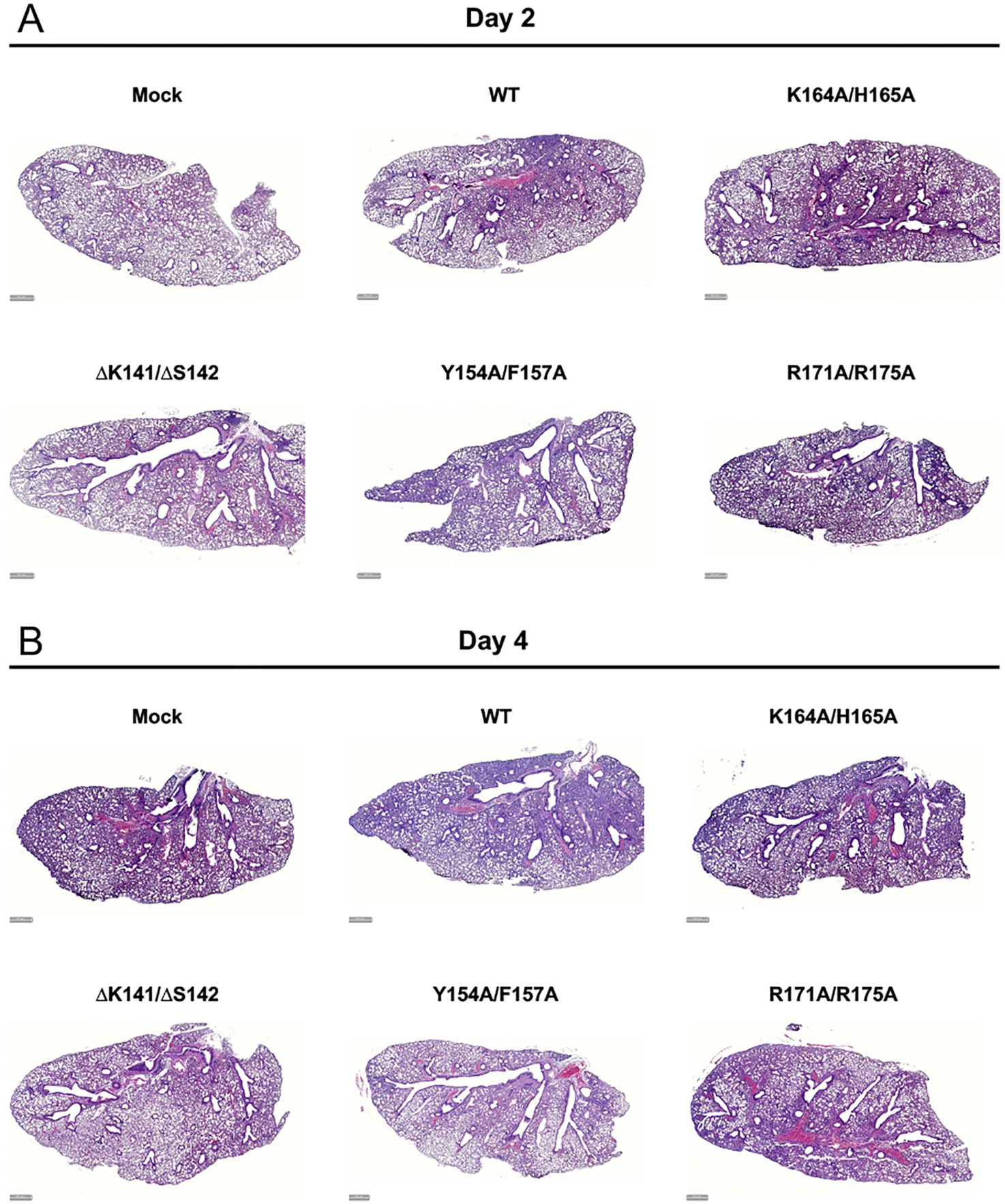
Lung histopathology of rSARS-CoV-2 Nsp1 mutant– infected mice. Whole-slide images of H&E-stained lung sections representative of each group at day 2 (**A**) and day 4 (**B**) post-infection (Scale bar = 500µM).

**Supplemental Figure 3:**
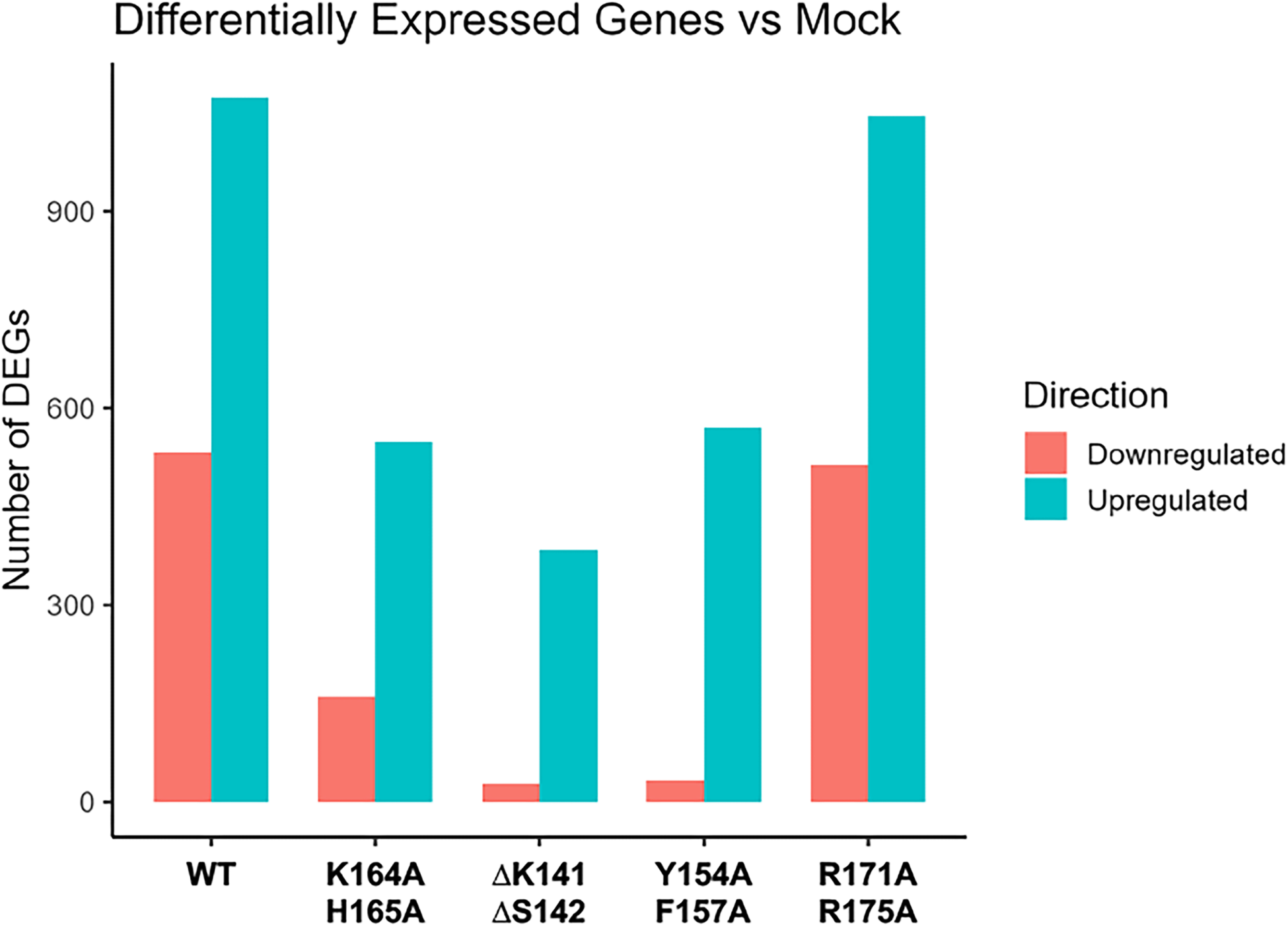
Lung differentially expressed gene (DEG) following rSARS-CoV-2 infection. Lung tissues collected from mice infected with rSARS-CoV-2 WT and Nsp1 mutants at day 4 post-infection were homogenized in TRIzol for RNA isolation and subjected to RNA sequencing (n = 3 per group). Differential expression analysis in WT-and Nsp1 mutant–infected mice were analyzed relative to mock– infected mice and was performed using a threshold of adjusted *p* value (padj) < 0.05 and absolute log2 fold change ≥ 1. Downregulated genes are shown in red and upregulated genes in blue.

## REFERENCES

1. Drosten C, Günther S, Preiser W, Van Der Werf S, Brodt H-R, Becker S, et al. Identification of a novel coronavirus in patients with severe acute respiratory syndrome. New England journal of medicine. 2003;348(20):1967–76.

2. Tg K. A novel coronavirus associated with severe acute respiratory syndrome. N Engl J Med. 2003;348:1953–66.

3. Coleman CM, Frieman MB. Emergence of the Middle East respiratory syndrome coronavirus. PLoS pathogens. 2013;9(9):e1003595.

4. De Groot RJ, Baker SC, Baric RS, Brown CS, Drosten C, Enjuanes L, et al. Commentary: Middle east respiratory syndrome coronavirus (mers-cov): announcement of the coronavirus study group. Journal of virology. 2013;87(14):7790–2.

5. Zhou P, Yang X-L, Wang X-G, Hu B, Zhang L, Zhang W, et al. A pneumonia outbreak associated with a new coronavirus of probable bat origin. nature. 2020;579(7798):270–3.

6. Lu R, Zhao X, Li J, Niu P, Yang B, Wu H, et al. Genomic characterisation and epidemiology of 2019 novel coronavirus: implications for virus origins and receptor binding. The lancet. 2020;395(10224):565–74.

7. Wong L-YR, Perlman S. Immune dysregulation and immunopathology induced by SARS-CoV-2 and related coronaviruses—are we our own worst enemy? Nature Reviews Immunology. 2022;22(1):47–56.

8. Maurina SF, O’Sullivan JP, Sharma G, Rodriguez DCP, MacFadden A, Cendali F, et al. An evolutionarily conserved strategy for ribosome binding and host translation inhibition by β-coronavirus non-structural protein 1. Journal of molecular biology. 2023;435(20):168259.

9. Schubert K, Karousis ED, Ban I, Lapointe CP, Leibundgut M, Bäumlin E, et al. Universal features of Nsp1-mediated translational shutdown by coronaviruses. Molecular Cell. 2023;83(19):3546–57. e8.

10. Parenti NA, Cusic R, Renner DM, Jackson N, Ye C, Tan LH, et al. SARS-CoV-2 and MERS-CoV disrupt host protein synthesis via nsp1 with differential effects on the integrated stress response. Proceedings of the National Academy of Sciences. 2026;123(15):e2536296123.

11. Schubert K, Karousis ED, Jomaa A, Scaiola A, Echeverria B, Gurzeler L-A, et al. SARS-CoV-2 Nsp1 binds the ribosomal mRNA channel to inhibit translation. Nature structural & molecular biology. 2020;27(10):959–66.

12. Thoms M, Buschauer R, Ameismeier M, Koepke L, Denk T, Hirschenberger M, et al. Structural basis for translational shutdown and immune evasion by the Nsp1 protein of SARS-CoV-2. Science. 2020;369(6508):1249–55.

13. Mendez AS, Ly M, Gonzalez-Sanchez AM, Hartenian E, Ingolia NT, Cate JH, et al. The N-terminal domain of SARS-CoV-2 nsp1 plays key roles in suppression of cellular gene expression and preservation of viral gene expression. Cell Reports. 2021;37(3).

14. Zhang K, Miorin L, Makio T, Dehghan I, Gao S, Xie Y, et al. Nsp1 protein of SARS-CoV-2 disrupts the mRNA export machinery to inhibit host gene expression. Science Advances. 2021;7(6):eabe7386.

15. Mei M, Cupic A, Miorin L, Ye C, Cagatay T, Zhang K, et al. Inhibition of mRNA nuclear export promotes SARS-CoV-2 pathogenesis. Proceedings of the National Academy of Sciences. 2024;121(22):e2314166121.

16. Shehata SI, Parker R. SARS-CoV-2 Nsp1 mediated mRNA degradation requires mRNA interaction with the ribosome. RNA biology. 2023;20(1):444–56.

17. Li J, Wang K, Wang J, Zhong C, Wang S, Peng H, et al. SARS-CoV-2 nonstructural protein 1 suppresses host transcription by reducing RNA polymerase II levels. iScience. 2025;28(12).

18. Kamitani W, Narayanan K, Huang C, Lokugamage K, Ikegami T, Ito N, et al. Severe acute respiratory syndrome coronavirus nsp1 protein suppresses host gene expression by promoting host mRNA degradation. Proceedings of the National Academy of Sciences. 2006;103(34):12885–90.

19. Zhou F, Periasamy S, Jackson ND, Cheng WS, Soto Acosta R, Tripathi A, et al. Redundant and distinct mechanisms suppress innate immune activation during SARS-CoV-2 infection. PLoS Biol. 2026;24(5):e3003808.

20. Liu S, Stauft CB, Selvaraj P, Chandrasekaran P, D’Agnillo F, Chou C-K, et al. Intranasal delivery of a rationally attenuated SARS-CoV-2 is immunogenic and protective in Syrian hamsters. Nature Communications. 2022;13(1):6792.

21. Benedetti F, Snyder GA, Giovanetti M, Angeletti S, Gallo RC, Ciccozzi M, et al. Emerging of a SARS-CoV-2 viral strain with a deletion in nsp1. Journal of Translational Medicine. 2020;18(1):329.

22. Chen Y, Wang J, Liu C, Su L, Zhang D, Fan J, et al. IP-10 and MCP-1 as biomarkers associated with disease severity of COVID-19. Molecular Medicine. 2020;26(1):97.

23. Polese B, Ernst M, Henket M, Ernst B, Winandy M, Njock M-S, et al. Circulating inflammatory cytokines predict severity disease in hospitalized COVID-19 patients: A prospective multicenter study of the European DRAGON consortium. Journal of infection and public health. 2024;17(12):102589.

24. Ranjbar M, Rahimi A, Baghernejadan Z, Ghorbani A, Khorramdelazad H. Role of CCL2/CCR2 axis in the pathogenesis of COVID-19 and possible Treatments: All options on the Table. International immunopharmacology. 2022;113:109325.

25. Ye C, Chiem K, Park J-G, Oladunni F, Platt RN, Anderson T, et al. Rescue of SARS-CoV-2 from a single bacterial artificial chromosome. MBio. 2020;11(5):10.1128/mbio.02168-20.

26. Gen R, Addetia A, Asarnow D, Park Y-J, Quispe J, Chan MC, et al. SARS-CoV-2 nsp1 mediates broad inhibition of translation in mammals. Cell Reports. 2025;44(5).

27. Ye C, Ezzatpour S, Imbiakha B, Jackson N, Cagatay T, Cupic A, et al. Deletion of a KSF motif attenuates NSP1 host cell translation shutoff and impairs SARS-CoV-2 virulence. bioRxiv. 2025:2025.04. 16.649178.

28. Vora SM, Fontana P, Mao T, Leger V, Zhang Y, Fu T-M, et al. Targeting stem-loop 1 of the SARS-CoV-2 5′ UTR to suppress viral translation and Nsp1 evasion. Proceedings of the National Academy of Sciences. 2022;119(9):e2117198119.

29. Liao M, Liu Y, Yuan J, Wen Y, Xu G, Zhao J, et al. Single-cell landscape of bronchoalveolar immune cells in patients with COVID-19. Nature medicine. 2020;26(6):842–4.

30. Wauters E, Van Mol P, Garg AD, Jansen S, Van Herck Y, Vanderbeke L, et al. Discriminating mild from critical COVID-19 by innate and adaptive immune single-cell profiling of bronchoalveolar lavages. Cell research. 2021;31(3):272–90.

31. Ferreira HAS, Guimarães LC, Costa PAC, de Campos CLV, Júnior SRAS, Queiroz-Júnior CM, et al. Targeting CCL2 silencing using polymer-lipid hybrid nanoparticles to reduce cytokine storm and inflammation in severe COVID-19. International Journal of Pharmaceutics. 2025:126073.

32. Channappanavar R, Perlman S, editors. Pathogenic human coronavirus infections: causes and consequences of cytokine storm and immunopathology. Seminars in immunopathology; 2017: Springer.

33. Trujillo JA, Fleming E, Perlman S. Transgenic CCL2 Expression in the Central Nervous System Results in a Dysregulated Immune Response and Enhanced Lethality after Coronavirus Infection. Journal of Virology. 2012;87:2376-89.

